# From Brownian to deterministic motor movement in a DNA-based molecular rotor

**DOI:** 10.1101/2024.01.26.577361

**Authors:** Florian Rothfischer, Matthias Vogt, Enzo Kopperger, Ulrich Gerland, Friedrich C. Simmel

## Abstract

Molecular devices that have an anisotropic, periodic potential landscape can be operated as Brownian motors. When the potential landscape is cyclically switched with a chemical reaction or an external force, such devices can harness random Brownian fluctuations to generate directed motion. Recently, directed Brownian motor-like rotatory movement was demonstrated with an electrically switched DNA origami rotor with designed, ratchet-like obstacles. Here, we demonstrate that also the intrinsic anisotropy of DNA origami rotors that originally were not designed as Brownian motor devices is sufficient to result in motor movement. We show that for low amplitudes of an external switching field such devices operate as Brownian motors, while at higher amplitudes the movement is better described by the deterministic motion of an overdamped electrical motor. We characterize the amplitude and frequency dependence of the movements in both regimes, showing that after an initial steep rise the angular speed peaks and drops for excessive driving amplitudes and frequencies. The characteristics of the rotor movement are well described by a simple stochastic model of the system.

## I. INTRODUCTION

One of the major differences between nanoscale devices and macroscopic machines is the dominance of Brownian motion at small scales and thus the presence of large thermal fluctuations that are superimposed on the desired movement of the devices. Biology has evolved intricate mechanisms for the operation of its molecular machines that allow the ‘rectification’ of undirected Brownian movement, and thus enable locomotion and the generation of forces at the nanoscale. An important abstraction of the underlying mechanism of these machines is the concept of the Brownian ratchet [1–9]. Ratchet systems have a periodic, but anisotropic potential landscape (i.e., potentials with a broken symmetry), which can be switched by an external field or a chemical reaction. In biology, ratchet-like mechanisms are thought to be involved in the movement of DNA and RNA polymerases [10–13], the synthesis of ATP by F0-F1 ATPase [14, 15], and linear transport motors such as kinesin, even though their full stepping mechanism is likely more complicated [16].

There have been various attempts to implement synthetic Brownian motors in physical and chemical systems. Directed movement was demonstrated using microparticles optically trapped in a ratchet potential [17, 18], in microfluidic systems [19], and also in synthetic molecular motors driven by light [20]. More recently, the DNA origami technique [21–23] was utilized to create a DNA-based rotary device with explicitly designed ratchet-like obstacles [24]. In this study, the origami rotor was driven out of equilibrium using an electrical field, whose direction was periodically switched from one direction to its opposite (i.e., by 180°), and back. A similar mechanism had been previously studied theoretically in Refs. [25, 26]. Notably, the rotor displayed directional rotation, with the direction of movement determined by the relative orientation of the electric field and the origami structure. Importantly, the irrotational field by itself would not lead to any biased rotor movement, but its superimposition with the potential of the origami structure does.

In a different study, we had previously characterized the intrinsic potential landscapes of various DNA origami rotor structures, in which a DNA rotor arm was attached to a pivot point on a DNA origami base plate via single-stranded DNA connectors [27]. In the absence of an external field, these origami rotors almost always displayed two preferred orientations, inadvertently resulting in a rotatory energy landscape with two minima. We surmised that, similarly as in Ref. [24], an externally applied irrotational electric field that is misaligned with the intrinsic potential minima would create an anisotropic effective energy landscape, which would allow the device to convert Brownian motion into directed rotatory motion when driven out of equilibrium.

In the following, we demonstrate that when periodically switching the DNA origami rotors with sufficiently low external fields, the device indeed behaves like a Brownian motor. Notably, our device also allows the application of much higher electric fields than those used in the previous study [24]. For high fields, the device transitions from the Brownian to a deterministic regime, where its steps are clocked by the external frequency, and where it behaves like an ‘overdamped electromotor.’ At higher frequencies as well as for very high driving fields, the rotor velocity is reduced, which is in agreement with an idealized theoretical model of the system.

## II. RESULTS

### A. Design and characterization of the DNA origami rotors

Our rotor structure [27, 28] consists of a 55 nm x 55 nm stator base plate with a DNA six-helix bundle (6HB) - the rotor arm - attached to the center of the plate (Figure 1a). The stator is immobilized on the substrate via biotin-streptavidin linkages, while the 463 nm long rotor arm is connected to the stator via a flexible joint. The rotor arm itself consists of two subunits, the first of which has a length of 50 nm and includes the joint and connection to the stator. The second subunit comprises a 6HB with a length of 413 nm and is attached to the first subunit as an extension. In previous work [28], the joint connecting the arm to the plate consisted of two single-strands of DNA, which were shown to wind around each other during rotation of the arm until it stalled. To facilitate unlimited unidirectional rotation, we enzymatically cut one of the connecting strands as indicated in the Figure. Figure 1b shows AFM and TEM images of the resulting DNA nanostructure.

**FIG. 1.**
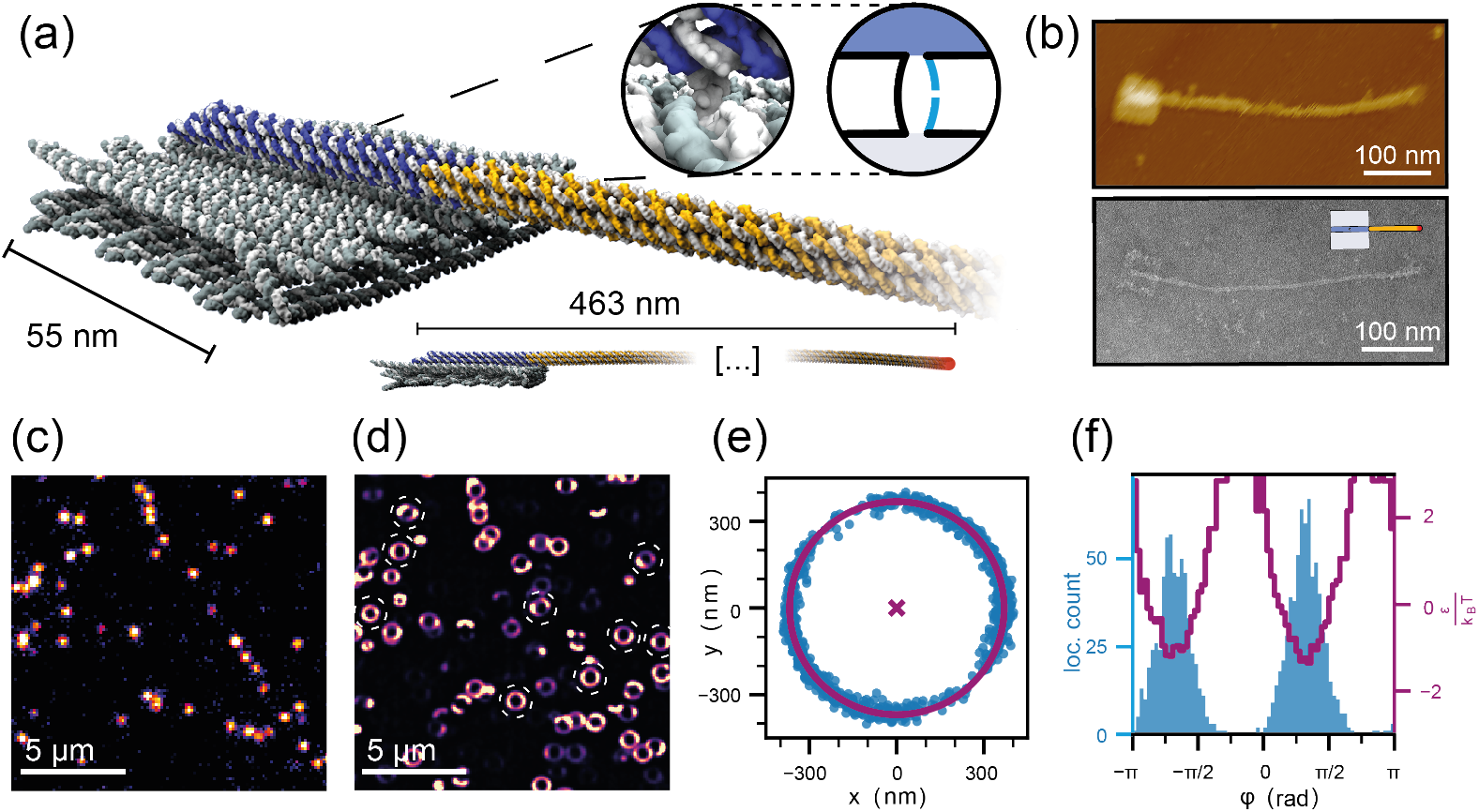
Structure design and data analysis workflow. (a) DNA origami nanostructure consisting of a 55 nm x 55 nm stator base plate, which bears a 463 nm long rotor arm connected to it via a flexible joint. The rotor consists of two subunits. A 50 nm long part (blue) connected to the stator through the joint and a 413 nm long rotor extension (yellow). The rotor is labelled at the tip with 42 fluorescent dyes (indicated by the red spot). The flexible joint consists of two single stranded DNA domains of the scaffold, one of which is cut to enable free rotation (cf. magnified inset). (b) AFM (top) and TEM (bottom) images of the fully assembled nanostructure. In schematic images we use the pictogram shown in the inset for the structure. (c) Single camera frame of the acquired video data. The fluorescence intensity is displayed in pseudo-color. (d) Pseudo-color localization heatmap of the total acquired data of one experiment. Dashed circles show freely rotating structures that are selected for further analysis. (e) Fitting a circle to the tracking data of a single structure allows to determine the position of the stator plate in the center and transform positional coordinates of individual localizations into relative polar coordinates. (f) The angular positions explored by the arm via diffusion (blue histogram) allow the calculation of the intrinsic energy landscape of our structure (shown in red), which exhibits two energy minima separated by *≈* 180°.

To track the motion of the rotors, we labelled the tips of the arms with 42 dye molecules each and recorded fluorescence microscopy videos in a total internal reflection fluorescence microscopy (TIRFM) setup. Figure 1c shows an example area of a single camera frame (exposure time 5 ms) containing several rotor structures. The movement was slowed down to reduce motion blur by increasing the viscosity of the buffer with 48% (w/w) sucrose. Computer-controlled electric actuation protocols were synchronized with the camera acquisition. The position of each rotor structure was localized with nanometer precision using the Picasso software package [29]. Figure 1d shows a localization heat map of the acquired video data for a single experiment. Structures that show free rotation and thus appear to have a correctly cut, single-stranded joint are selected for further analysis (dashed circles in Figure 1d). The center of each structure is determined with a circular fit to the fluorescence data collected from it (Figure 1e), and the positional coordinates of all localizations are subsequently transformed into relative polar coordinates. The rotor orientation with respect to the external camera reference frame is denoted as *φ*, while the integrated angular distance covered by each rotor over the course of the measurement is defined as *φ*_*c*_. Experimental details and a description of actuation protocol and data analysis are given in the Supplemental Material [30].

As demonstrated previously [27, 31], the rotors assume two preferred orientations in free diffusion measurements (Figure 1f). Histograms of the rotor positions provide an estimate for the equilibrium angle distribution *p*(*φ*), which in turn allows us to reconstruct the intrinsic potential energy landscape *E*_*in*_ of the rotor on top of the stator by inverting the Boltzmann relation, i.e., *E*_*in*_(*φ*) = −*k*_*B*_*T* ln [*p*(*φ*)]. The resulting potential can be well approximated by the function

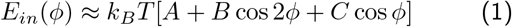

with the fit parameters *A* = 0.98, *B* = −2.24, *C* = −0.108. This function is 2*π*-periodic and has minima at *ϕ* = 0 and *ϕ* = *π*. We arbitrarily defined the angle of the lower minimum as *ϕ* = 0, which we take as the reference angle of the rotor structure (the angle *ϕ* is measured with respect to this angle, while *φ* is measured with respect to the camera frame).

### B. Brownian motor movement at low external driving

We experimentally assessed whether the DNA rotors would display directional movement as expected (Figure 2a) by monitoring a large number of individual rotors randomly oriented on a microscopic cover slide. To switch the potential, we applied a voltage protocol *V* (*t*) that periodically generated 100 ms long electrical pulses with a given field direction, followed by 100 ms long pulses in the opposite direction (i.e., *V* (*t*) = *V*_0_ sgn(sin *ω*_0_*t*) with *f*_0_ = *ω*_0_*/*2*π* = 1*/*(200 ms) = 5 Hz). We tracked the movement of the tips of individual rotor arms for different values of *V*_0_ (170-400 structures per measurement) and determined the integrated angular displacement *φ*_*c*_(*t*) covered by the arms from the data. These were then used to calculate the effective angular velocities *ω*(*t*) of the rotor arms. We also determined the orientation of the individual rotor arm platforms with respect to the external field, characterized by the angular mismatch *α*, and then plotted the angular velocity of each arm as a function of this angle. Application of voltages below *V*_0_ = 20 V did not result in noticeable directional movement of the rotors (Figure 2b), which indicates that the intrinsic energy barriers were not sufficiently modulated by the external field. Directional movement was clearly observed, however, when applying higher voltage amplitudes (Figure 2c & 2d). The speed of the rotors had a sinusoidal dependence on the angular mismatch (αsin 2*α*), with a maximum absolute value at *α* = ± *π/*4, and vanishing net movement for *α* = 0, *±π/*2. As indicated, we find movement in the counter-clockwise (CCW) direction 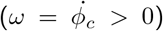 for 0 ≤ *α < π/*2, and clockwise (CW) rotation (*ω <* 0) for −*π/*2 ≤ *α <* 0.

**FIG. 2.**
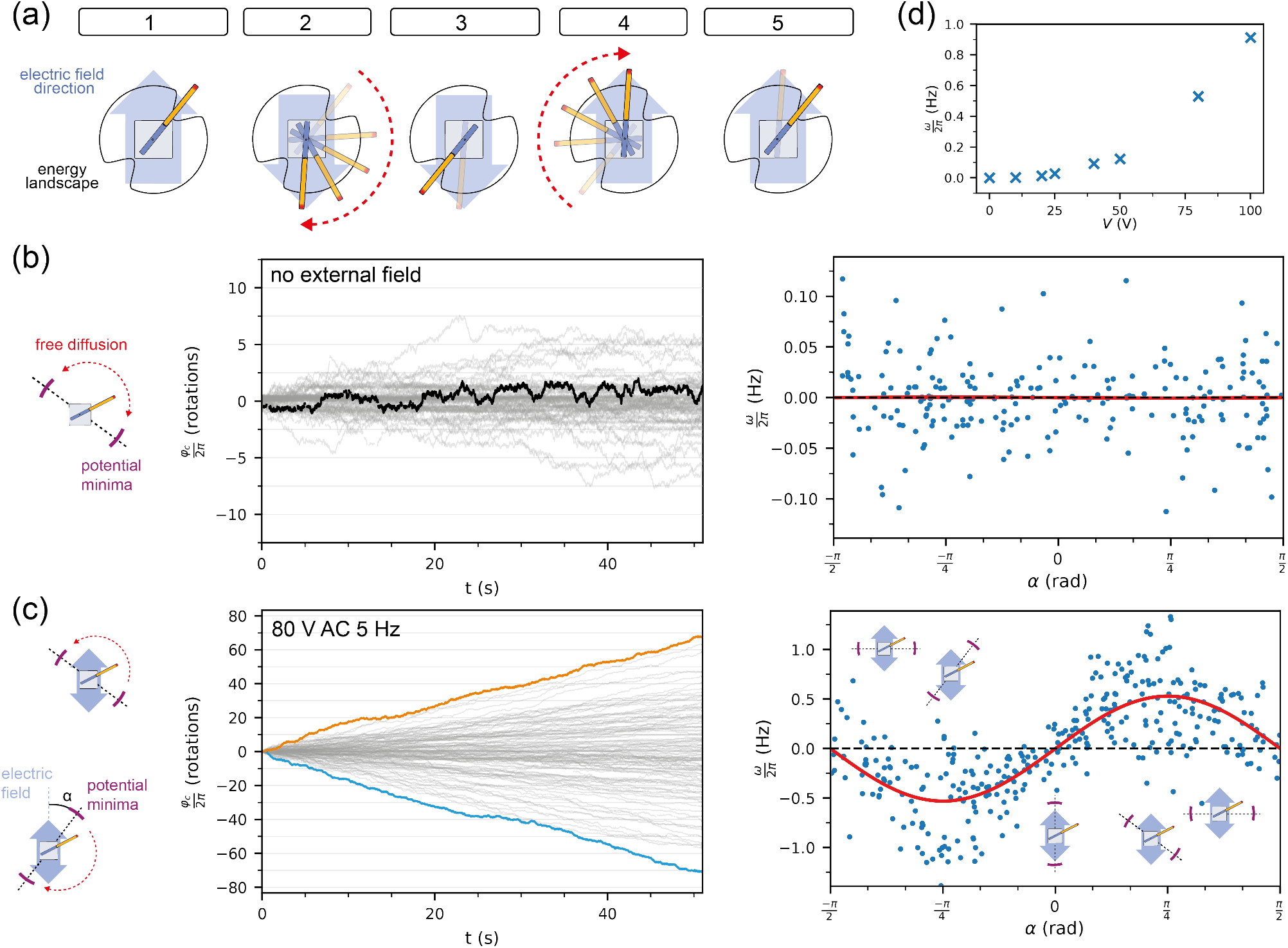
Brownian motor movement at low external driving. (a) Schematic explaining how the superposition of the time-dependent electric field and the intrinsic energy landscape can lead to directed movement. The field stabilizes the rotor arm in one of the two energy minima of the stator. When the field direction is switched, the arm preferentially moves in the clockwise direction for the particular arrangement shown here. (b) Left: Particle tracking traces recorded in the absence of an external field. The ensemble of all trajectories is shown in grey, an exemplary trace is highlighted in black. Right: The effective angular velocity *ω* for each trace is plotted against the corresponding angular mismatch α. (c) Left: Exemplary rotor arm traces in the presence of an applied external field (*V*0 = 80 V) whose direction is switched back and forth with *f*0 = 5 Hz. The blue trace shows a rotor moving predominantly in clockwise direction, while the orange trace displays anti-clockwise rotation. The full ensemble of trajectories is shown in transparent grey. The angle α is defined as the orientation of the closest intrinsic potential minimum with respect to the direction of the external field, i.e., α *>* 0 in the upper scheme (CCW rotation), and α *<* 0 (CW rotation) in the lower scheme. Right: Effective angular velocity *ω* for each trace on the left plotted against the corresponding angular mismatch α. *ω* shows a sinusoidal dependence on α on average. The red curve is a fit to the data with *ω*(α) = *ω*_*fit*_ sin 2α. The maximum angular speed (*±ω*_*fit*_) is observed for α = ±45° (d) Plot of *ω*_*fit*_ as a function of externally applied voltage.

### C. Angle dependence of the energy landscape

Our experimental observations can be rationalized by considering the superposition of the externally applied bias potential with the intrinsic potential landscape of the rotor *E*_*in*_. In thermodynamic equilibrium, the rotor arm diffusively moves in its intrinsic potential without directional bias, i.e., clockwise and counterclockwise movements cancel each other on average. Applying an alternating (AC) electric field with orientation −*α* with respect to the rotor platform will see-saw the intrinsic potential by adding an oscillatory term α*V*_0_ cos (*ϕ* + *α*) ×sgn(sin *ω*_0_*t*) (the argument of the cosine is *ϕ*+*α* due to the definition of the angle *α*, cf. Fig. 2). This results in a total effective potential that is given by

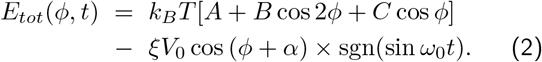

Here *ξ* represents the coupling strength that converts externally applied voltage into mechanical torsion energy. Importantly, the external field alone does not impose any directional movement on the rotor. However, the combined time-dependent potential *E*_*tot*_ can give rise to a ratchet-like effect whose strength depends on the angular mismatch *α*.

This can be easily understood by considering the shape of the combined potential in its two states for different angles *α* (Figure 3a). When the external field is in its positive or negative half-cycle, respectively, the total potential will have the shape

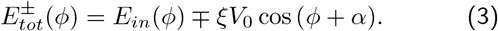

The intrinsic potential has a minimum at *ϕ ≈* 0 and, for small values of the parameter *C*, two neighboring maxima at *ϕ ≈ ±* π*/*2, which separate it from the other minimum at *ϕ ≈ ± π*. When the angular mismatch is *α* = 0 (more generally, *α* = *n · π* (*n ϵ* ℤ)), the external field pulls in the direction of one minimum in one half-cycle, and in the direction of the other minimum in the other half-cycle. Depending on the magnitude of the modulation *V*_0_, the arm will either remain in the local minimum, or transition to the other minimum. As the landscape is (almost) symmetric for *α* = 0, there is no directional bias for this transition. By contrast, for other angles *α*, one of the barriers will be reduced and the other elevated in each half-cycle of the electric field, favoring transitions between the minima to occur always in the same direction (see Figure 3a, cf. [5]). When the external modulation is strong enough, the rotor arm can escape the minimum with the help of thermal fluctuations, leading to CCW Brownian motor-like movement for 0 *< α < π/*2, and CW movement for −*π/*2 *< α <* 0. A stochastic simulation of the Langevin equation for the rotor system shows the expected behavior (cf. Supplemental Material Fig. S5 [30]). We find that the rectification effect is strongest for *α* = ± *π/*4, which is in agreement with our experimental observations (Supplementary Fig. S6 [30]).

**FIG. 3.**
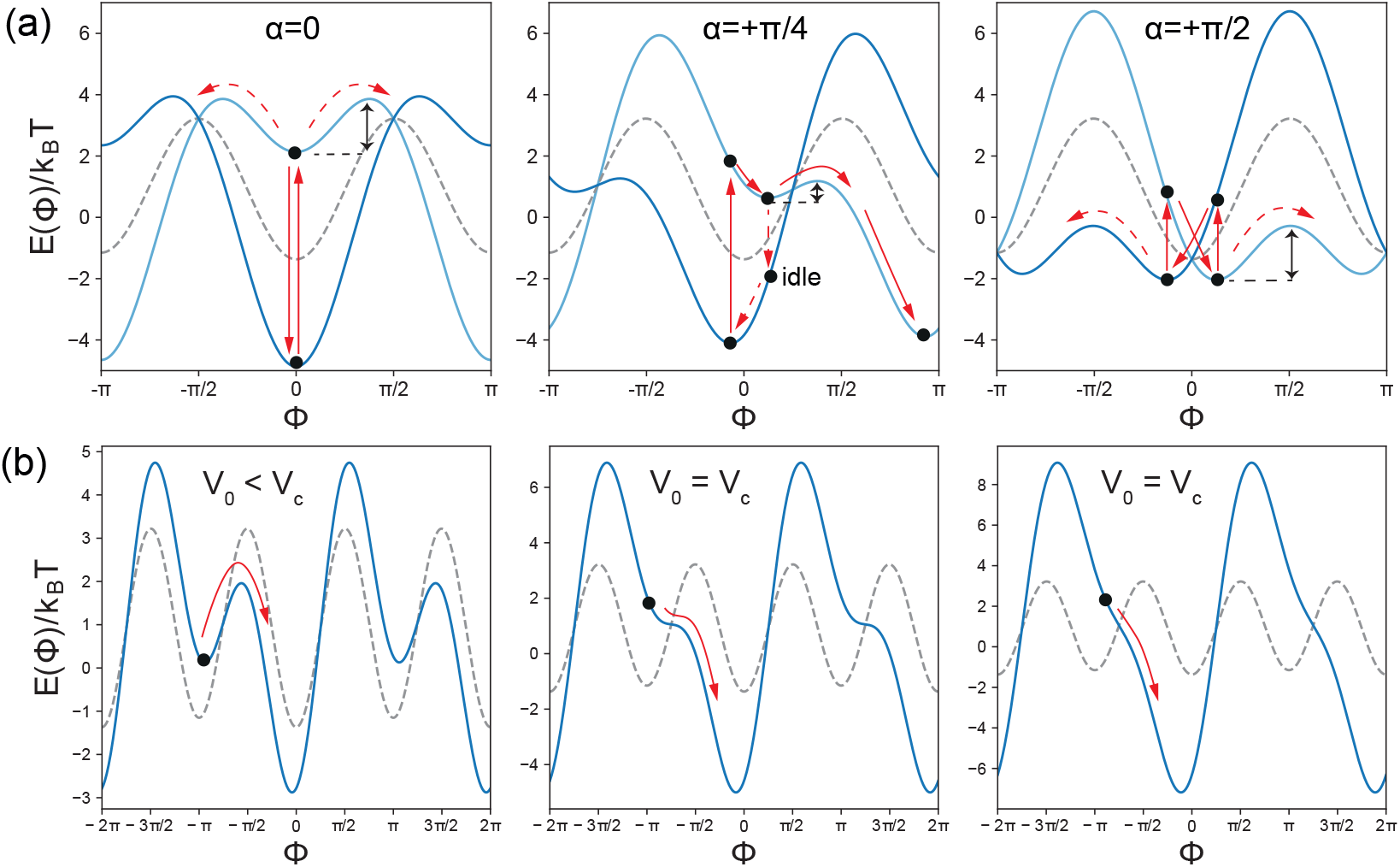
(a) Total potential as a function of Φ for different angular mismatches *α* (cf. Supplemental Material Fig. S3-S4 [30]). The dark blue curves show *E*_*tot*_(Φ) for an external potential of *ξV*_0_ = +3.5 *k*_*B*_*T*, the light blue curves correspond to the opposite polarity (*ξV*_0_ = −3.5 *k*_*B*_*T*). The intrinsic potential is shown with a grey dashed line. When for *α* = 0 the arm initially is in the minimum at Φ = 0 (indicated by the black spot on the dark blue curve), switching the potential will lift it to the corresponding minimum at Φ = 0 of the light blue curve. As indicated by the red arrows, it may stay at Φ = 0, or escape either to the left (CW) or right (CCW) with a probability that depends on the corresponding energy barriers (one is indicated by a black double arrow). For *α* = +*π/*4 the barrier to the left is elevated, while the barrier to the right is reduced and the arm will preferentially transition to the minimum on the right after switching (leading to CCW movement). When it does not escape from the higher minimum on the light blue curve before the potential is switched again, it may undergo an idle cycle as indicated. For *α* = +*π/*2, transitions between the two states of the total potential also do not generate any net movement (for Brownian dynamics simulations in the potentials see the Supplemental Material Fig. S5 & S7 [30]). (b) Total potential for *α* = +*π/*4 for external voltages below, at, and above the critical value *V*_*c*_ = 4.55 *k*_*B*_*T/ξ* (grey dashed line: intrinsic potential). For *V*_0_ < *V*_*c*_, the potential has a low and high minima close to Φ = 0, 2 *π* and Φ = *π*. The rotor arm can thermally escape from the minima at Φ = ± *π*, leading to Brownian motor movement in CCW direction. At *V*_0_ = *Vc*, the minima close to Φ =±*π* become saddle points, and at *V*_0_ > *V*_*crit*_ the movement to the right is always downhill until the arm reaches one of the deep minima.

### D. Transition to quasi-deterministic motor movement

Our model further suggests that for high enough voltages the energy landscape will transition from a potential with two minima within [ −*π, π*] to a potential with only one minimum (Figure 3b). When for *α* = *π/*4 the external field is increased, the potential minima and maxima that initially were at *ϕ ≈ π* (*mod* 2*π*) and *ϕ ≈* 3*π/*2 (*mod* 2*π*), respectively, move towards each other and merge in a saddle point at *ϕ* = 5*π/*4 (*mod* 2*π*) when *V*_0_ *≈* 4.55 *k*_*B*_*T/ξ* (Figure 3b). Thus, for strong enough fields the energy barrier for movement in one direction actually vanishes and therefore the rotor arm can move without assistance of thermal fluctuations. In this regime, the overdamped system is expected not to behave as a Brownian motor, but to move quasi-deterministically within its potential landscape (‘quasi-’ as its movement is still subject to thermal fluctuations).

As our simple rotor design allows application of much higher voltages than the more complex multicomponent origami rotors studied previously [24], we set out to also explore the predicted deterministic regime. As shown in Figure 4, the rotor moves much faster when rectangular voltage signals with amplitudes above ≈ 100 V are applied. Furthermore, the *φ*_*c*_(*t*) traces indicate that, in contrast to the Brownian motor regime, the rotor indeed rarely reverts its direction (Figure 4a). As anticipated, also in the deterministic case the rotor velocity has a sinusoidal dependence *ω* ∝ sin 2*α* and the movement of the rotor arms is observed fastest for *α* = *π/*4 (Figure 4b).

**FIG. 4.**
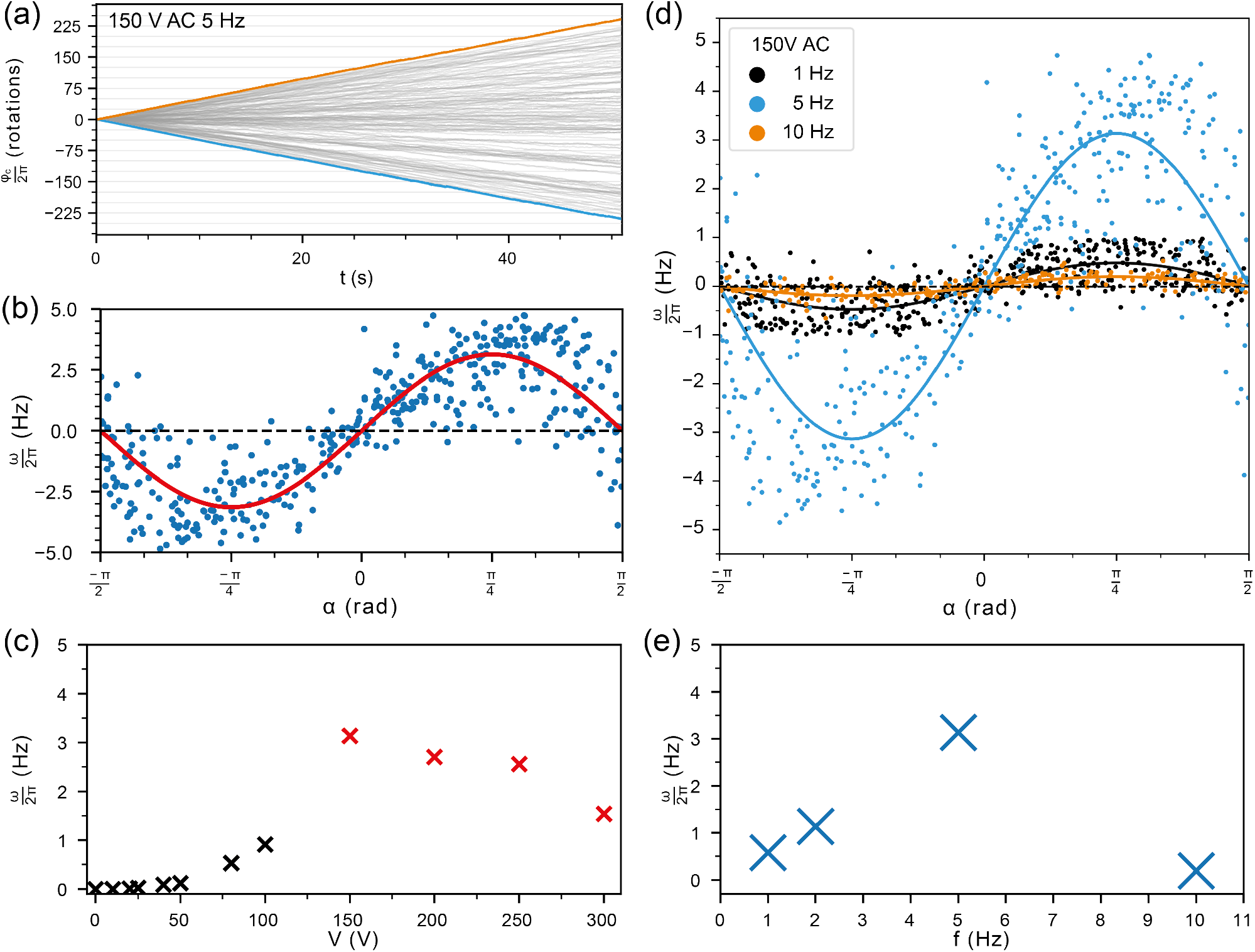
Rotor movement at higher driving voltages. (a) Exemplary traces at 150 V applied external AC field at *f*0 = 5 Hz. The blue trace shows deterministic movement in a clockwise direction, while the orange trace depicts a structure rotating counter-clockwise. The full ensemble rotor trajectories is shown in transparent grey. (b) Effective angular velocity *ω* vs. angular mismatch *α* for *V*_0_ = 150 V. The red curve is a fit to the data with the function *ω*(*α*) = *ω*_*fit*_ sin 2*α*. (c) Plot showing the dependence of *ω*_*fit*_ on the applied voltage, combining low voltage (cf. Fig. 2d) and high voltage data. (d) Frequency dependence of deterministic rotor movement at *V*_0_ = 150 V, displaying *ω* as a function of *α* for measurements with *f*_0_ = 1 Hz (black), 5 Hz (green), and 10 Hz (red), and corresponding sine fits. (e) Plot of the angular velocity *ω*_*fit*_ for all measured frequencies.

For *f*_0_ = 5 Hz, we observed the highest speed with a rectangular signal around *V*_0_ = 150 V, which resulted in a rotation frequency of *f* = *ω/*2*π ≈* 3.1 Hz. Notably, application of higher voltages leads to a reduction in the frequency of the rotor rotation, e.g., to *f ≈* 1.6 Hz at *V*_0_ = 300 V (Figure 4c). Our model for *E*_*tot*_(*ϕ, t*) (Eq. (3)) indicates that for higher voltages the potential landscape is completely dominated by the externally imposed potential and thus becomes almost symmetric. Stochastic simulations show that under these conditions the rectification effect observed for the lower voltages is indeed diminished (cf. Supplemental Material Fig. S8 [30]).

As the maximum possible angular velocity is given by *f*_0_ = 5 Hz, we only achieve *ω*_*max,exp*_ ≈ 0.61 ×*ω*_*max,theo*_ at this frequency. This suggests that the rotor arm, while not rotating back, is not able to follow the external field in every cycle (i.e., it has ≈39% idle cycles). In order to study the frequency dependence of the rotor arm movement in more detail, we systematically changed the switching fre-quency *f*_0_ for a constant voltage amplitude of *V*_0_ = 150 V (Figure 4d & e). While the speed steadily rises when going from *f*_0_ = 1 Hz to 2 Hz and finally 5 Hz, it is strongly reduced for *f*_0_ = 10 Hz. This behavior is a consequence of the overdamped movement of the rotor arm. For higher frequencies, the rotor cannot follow the switching of the potential landscape any more, and therefore increasingly undergoes idle cycles with Δ*ϕ* = 0. The critical frequency *f*_*c*_ is thus set by the friction coefficient *γ*_*r*_ of the rotor arm, which is determined by its geometry and the viscosity of the medium. The frequency dependence of the rotor arm movement can be recapitulated in Brownian dynamics simulations (Supplemental Material Fig. S9-S11 [30]), which show that for large enough voltages *f*_*c*_ ∼ *V*_0_*/γ*_*r*_.

## III. DISCUSSION

We have shown that a rotary DNA origami nanodevice ‘serendipitously’ acts as a Brownian motor due to its intrinsic energy landscape that contains two energy minima with a depth on the order of 1 *k*_*B*_*T*. These energy minima were not explicitly designed, but are an emerging feature of the structure. In previous work [27] we found it challenging to create nanomechanical origami structures with a completely flat energy landscape, and a modulation of the potential on the order of *k*_*B*_*T* is rather typical. In the specific case of our rotor structures this means that the rotor arm has two preferential orientations with respect to the stator base plate. When the rotor arm diffusively explores its mechanical energy landscape, it will randomly transition between these minima without any directional bias, leading to overall zero net movement.

Directional movement of the rotor arm can be induced by externally applying an electric field that is switched back and forth between two opposite directions. In such a setting the electric field alone does not provide any directional bias, but the superimposed potential generated by the intrinsic mechanical landscape and the external field does. Depending on the relative orientation of the external field and the intrinsic minima (measured by the angular mismatch *α*), the effective potential landscape will be more or less asymmetric. When the system is driven out of equilibrium by switching between two alternative asymmetric potentials (corresponding to the two field directions), the diffusive movement of the rotor arm can be rectified. The effect turns out to be maximal for *α* = ±*π/*4.

The overall behavior of the system in the low voltage regime is reminiscent of a Brownian ratchet [5]. The canonical flashing ratchet model usually considers an intrinsically asymmetric energy landscape that can be modulated or switched completely on and off. In our setting the asymmetry is created by the superposition of the external and intrinsic potential. Even though our measured intrinsic potential is itself slightly asymmetric (due to the non-vanishing term *C* in Eq. 1), simulations indicate that this is not a necessary requirement. As a caveat, we have to note that our estimated intrinsic potential is only an approximation. First, the Boltzmann inversion is likely not accurate for angular positions close to the potential barriers, where we naturally cannot collect a large number of data points. Furthermore, we do not know whether the intrinsic landscape will be changed in the presence of electric fields (e.g., via deformation of the structures).

What is notable from a nano-engineering perspective is that the Brownian motor property of our rotors essentially comes for free. This suggests that it is actually relatively easy to make such devices as we would always expect some energetic ‘corrugation’ of the mechanical energy landscape of such constructs. The intrinsic motor characteristics of the structures can then serve as a starting point, guiding rational design choices towards enhanced motor performance. It is maybe not too far-fetched to draw a comparison to the evolution of membrane-embedded molecular rotors in biology [32]. It is conceivable that ancestral membrane-bound protein assemblies acting as stators and rotors had evolved early and naturally showed some directional movement in the presence of a transmembrane gradient simply due to their serendipitous asymmetry. This movement could then have been further improved through evolutionary optimization [32, 33].

For low amplitudes of the external field, thermal fluctuations are required to drive the movement - and in this sense the system indeed is a Brownian motor whose mechanism allows to rectify the otherwise undirected thermal movement. Accordingly, in our Langevin simulations the movement of the rotor arm ceases when the fluctuation term is set to zero (Supplemental Fig. S5 [30]). Notably the behavior is different for the high voltage case, where the intrinsic landscape is distorted by the external potential to such a degree that the rotor arm always moves energetically down-hill after switching the potential, and thermal fluctuations are not required for the movement (Supplemental Fig. S4& S7 [30]).

With an estimated rotational drag coefficient of *γ*_*r*_ = 1.1 pN nm s [28, 30] and a moment of inertia of *I* = *ML*^2^*/*3 = 6 10–^34^kg m^2^ that can be derived from the mass *M* of the ≈ 8000 DNA base-pairs (with 650 Da per bp) comprising the arm and its length of *L* ≈ 463 nm, the relaxation time *t*_*r*_ = *I/γ*_*r*_ is way below 1 ps. The rotor arm thus is completely overdamped and in the quasi-deterministic regime will move with a angular drift velocity that is given by *ω*_*drift*_ = *τ/γ*_*r*_ = −1*/γ* · *dE*_*tot*_*/dϕ*. Due to the absence of inertia, the rotor arm will immediately stop when the field is switched off.

It is instructive to compare the movement of our over-damped motor to a simple macroscopic two-pole DC electromotor made from a rotating coil, a permanent magnet, and a commutator. In this case the commutator ensures periodic switching of the direction of the electric current through the coils, changing the magnetization of the rotor and thus driving it through its cycle. Inertia is crucial for such a motor as it ensures that the rotor will continue to rotate in the same direction and does not stall when the brushes lose contact with the commutator. Such a motor would not work in the overdamped regime.

In conclusion, we have shown that a rotary DNA origami nanodevice comprised of a DNA rotor arm attached on a rigid stator plate can be repurposed as a nanoscale electromotor by externally applying a non-rotating, periodically switched electrical field whose direction is mismatched with the minima of its intrinsic energy landscape. Depending on the strength of the electric field and the corresponding modulation of the potential energy, the rotor acts as a Brownian motor - utilizing and rectifying the thermal fluctuations of the system -, or as an overdamped electrical motor. In particular, our results show that a Brownian motor can be generated without explicit design, but simply by utilizing unavoidable irregularities in the potential landscape of a DNA-based nanodevice.

We have here used an external electric field to drive the rotor out of equilibrium, as this approach is comparatively easy to implement experimentally. One of the major challenges for future work will be realization of out-of-equilibrium systems by other means, in particular Brownian motors that are fueled by chemical reactions [34, 35].

## ACKNOWLEDGMENTS

This work was funded by the Deutsche Forschungsgemeinschaft (DFG, German Research Foundation) through SFB 1032 (project ID 201269156 TPA2). We gratefully acknowledge funding by the BMBF through its 6G-life initiative. Florian Rothfischer has personally been funded by the Konrad-Adenauer-Stiftung (KAS). We thank the group of Prof. Hendrik Dietz for providing us with DNA origami scaffold strands. We thank Adrian Büchl for the TEM image. Source data and all other data that support the plots within this paper and other findings of this study are available from the corresponding author upon reasonable request. The source code of the data analysis routines and simulation files employed in this study are available from the corresponding author upon reasonable request.

