## Supplemental Information for "From Brownian to deterministic motor movement in a DNA-based molecular rotor"

Florian Rothfischer, Matthias Vogt, Enzo Kopperger, Ulrich Gerland, Friedrich C. Simmel*
*Department of Bioscience, TUM School of Natural Sciences,*
*Technical University Munich, D-85748 Garching, Germany*

(Dated: January 26, 2024)

**DNA origami preparation**

*Folding.* The DNA origami nanostructures (stator and rotor) were folded separately using in-house produced custom 8064 nt and 7704 nt scaffolds (both 100 nM in $ddH_2O$), kindly provided by the group of Prof. Hendrik Dietz. Linearization of the 8064 nt scaffold was performed using a restriction digest as described previously ([1]). All staple strands were ordered from Integrated DNA Technologies Inc. (100 µM in 10 mM Tris and 0.1 mM EDTA, pH 8.0, 1x TE). 100 µl of the scaffold was mixed with 3-fold excess of staples. Staple strands that contained a biotin modification or extensions for fluorescent dye attachment were added in 5-fold excess. The sample was adjusted to a final concentration of 1x TE and 20 mM $MgCl_2$. The DNA nanostructures were annealed in a thermal cycler (peqSTAR 2X; Peqlab Biotechnologie GmbH) over the course of 12 hours from $70°C$ to $40°C$. After folding, the structures were stored at $20°C$ until further processing.

*Purification.* Excess staples were removed by polyethylene glycol (PEG) precipitation of the origami structures similar as in previous work ([2]). Therefore, an equal volume of origami sample was mixed with PEG precipitation buffer (1x TE, 20 mM $MgCl_2$, 15% (w/v) PEG8k). After mixing the structures were centrifuged at 20,000 rcf at $20°C$ for 30 minutes. The supernatant was dropped and the pellet resuspended in 1x TE and 1 M NaCl. The purified stator structures were incubated with at least 20-fold molar excess of NeutrAvidin over biotin modified staples for 30 min at $20°C$. The purified rotor structures were incubated with 250-fold molar excess of ATTO 655 modified marker strands (ATTO655-atatgtacgggtctcactta). Subsequently, both structures were purified by agarose gel electrophoresis at 70 V for 1 h with a 0.5 % agarose gel using a gel buffer and running buffer containing 0.5× TBE, 12 mM $MgCl_2$ in a VWR Horizontal MINI S gel electrophoresis system (VWR International Ltd.).

*Stator-rotor assembly.* The final stator-rotor structures were assembled by incubation at $37°C$ for 30 min at 600 rpm. The rotor structures were added in 1.5-fold molar excess over the stator structures.

#### AFM imaging

AFM data was obtained using an Asylum Research Cypher ES (Oxford Instruments) AFM with Olympus BL-AC40TS-C2 cantilevers (Olympus) in AC mode. To prepare the sample, the structures were deposited on freshly cleaved mica by adding 5 µL of a 0.5 nM DNA origami solution (in $1\times$ TE and 20 mM $MgCl_2$) of the assembled nanostructure. The imaging buffer contained 95 µL of a $1\times$ TE and 200 mM $MgCl_2$ buffer with 15 µL of 620 mM $NiCl_2$ added for enhanced adhesion of the 6HB to the mica surface. Mixing of the imaging buffer on the chip was achieved by carefully pipetting up and down.

#### TEM imaging

The negative-stain TEM micrograph is recorded using the same experimental procedure as previously published by Büchl et al. [3] and Kopperger et al. [2].

#### Single-molecule TIRF microscopy and electrical actuation protocol

*Sample chambers.* The sample chambers are constructed as previously published [1] and consist of a custom made aluminium oxide ceramic top part (3 mm thickness, Laser-Cut-Processing GmbH, Germany) with buffer reservoirs, a PEG-Biotin modified cover slip and an interconnecting double sided tape layer (50 µm thickness, 3M 467MP transfer tape) which forms the flow channel. A custom 3D printed polypropylene electrode holder (Rapidobject GmbH, Germany) positions four 0.2 mm platinum wire electrodes in the buffer reservoirs.

*Electrical actuation setup for TIRFM.* Electrical actuation of our structures was performed by an in-house built electrical amplifier [1], controlled by the output of a DAQ-card. The DAQ cards output was controlled using a self-built Labview routine (source code available upon reasonable request) and consists either of a rectangular AC signal of 5 Hz with varying voltage (cf. Figure 2 and 4 of the main paper), or a rectangular AC signal with a voltage of 150 V and frequencies from 1 Hz to 10 Hz (cf. Figure 4).

*Conversion factor $\xi$.* The electrodes have a distance of 4.1 mm and therefore a voltage of $100\ \mathrm{V}$ translates into an electric field of $\mathcal{E} = 244\ \mathrm{V/cm}$. In earlier work, we obtained a conversion factor from electric field to exerted torque of $\kappa = 0.08\ \frac{\mathrm{pN\cdot nm}}{\mathrm{rad}}\frac{\mathrm{cm}}{\mathrm{V}}$ for the buffer conditions used here. Thus application of $100\ V$ would generate a maximum torque of $\tau = \kappa \times \mathcal{E} = 19.5\ \mathrm{pN\cdot nm}$. This directly translates into a maximum torsional potential energy $19.5\ \mathrm{pN} \cdot \mathrm{nm} = 4.75\ \mathrm{k_B T}$. Thus the conversion factor $\xi$ introduced in the paper would be $\xi = 0.0475\ k_B T/\mathrm{V}$. This conversion factor depends on various factors such as the quality of the surface treatment of the sample, however, and may vary between experiments. Our results suggest a higher value of $\xi \approx 0.1\ k_B T/\mathrm{V}$ for the experiments in the present work (see discussion below).

*PEG-biotin coated cover slips.* The microscopy slides (60x24 mm #1,5) from Menzel-Gläser (Germany) are produced in house as showed earlier in [2] by covalently attaching a PEG-Biotin (Biotin-PEG-silane MW 3,400, Laysan Bio Inc., USA) layer. For extended storage over several months in a freezer, we filled the storage container of the slides with Argon and drying beads.

*Sample chamber preparation for TIRF microscopy.* The sample chambers were prepared as previously published [1] with a flushing buffer containing 3 mM $MgCl_2$ and 30 mM TrisBorate and a high viscosity measurement buffer containing 3 mM $MgCl_2$, 30 mM TrisBorate and 48% (w/w) sucrose. 10 μl of the nanostructure sample (100 pM) was applied to the chamber for appropriate surface coverage.

*TIRF microscopy.* The custom-built objective-type single molecule TIRF microscopy setup is constructed as shown in [1]. It consists is based on an Olympus IX71 inverted microscope body (Olympus, Japan), a 642 nm excitation laser (Toptica iBeam smart, diode laser, 150 mW, Germany), a 100x oil immersion objective (UAPON 100xOTIRF objective, NA 1.49 oil, Olympus, Japan) and a filter cube equipped with a ZT532/640RPC dichroic mirror and a ZET532/640 (Chroma Technology, Germany) emission filter. All videos were acquired with an ORCA-Fusion Digital CMOS camera (Hamamatsu Photonics, Japan) with the sample chamber mounted on a U-780.DOS XY stage and a P-611.Z piezo z-stage (both Physik Instrumente (PI) GmbH, Germany).

**Data analysis**

The TIRFM videos were initially processed using the *Picasso* software package [4]. The combined emitter point spread functions of all fluorescent dyes at the labelled rotor tips were treated as a single emitter and therefore localized and fitted with a 2D Gaussian fit. If necessary, the drift correction functions of *Picasso* were applied to the data set. Particles that showed the expected movement (full rotations due to the cut joint) were picked manually and their event list was exported for subsequent analysis using a custom python-based routine. Localizations with a precision estimated to be worse than 50 nm (based on Picasso's "localization precision" parameter) were excluded from the analysis. The center of rotation of each particle was calculated by fitting a circle to its localizations. Subsequently, the positional coordinates of all localizations were transformed into relative polar coordinates. The cumulative particle traces were calculated by summing up the shortest angular difference between consecutive frame localizations. Our high time resolution (5 ms exposure) and the use of a viscous buffer (48 % (w/w) sucrose) ensure that angular differences between consecutive frames are sufficiently small ($\Delta\varphi_a \ll \pi$) and well resolved (cf. Supplemental Figure SS1). The relative positions of the energy landscape minima of each particle were determined by incorporating an initial measurement period with no electric field (5 s, not shown in the exemplary particle traces in the figures). The resulting free diffusion angular data was fitted with two Gaussian functions and their centers were extracted. The exemplary energy landscape shown in Figure 1F of the main paper was calculated as previously published in [3]. Particles displaying unusual behavior, such as no fluctuations due to Brownian motion or no response to the electric field, were discarded from the analysis as they were likely non-specifically bound to the substrate surface.

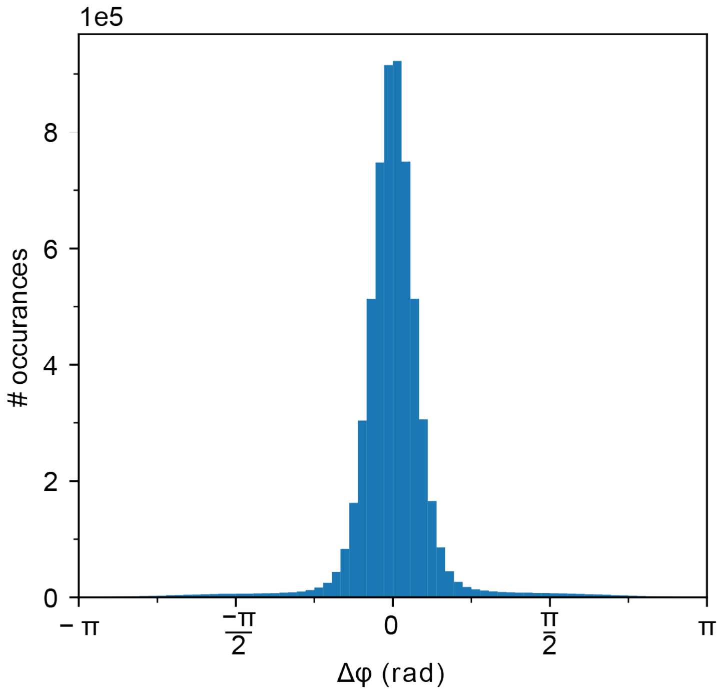


FIG. S1. **Histogram of angular differences between consecutive frames for all particles.** The angular difference is sufficiently small to reliably calculate a cumulative particle trace without missing parts of the particle movement.

**Structure design**

The stator-rotor nanostructure in this study is constructed from two separate DNA origami nanostructures. The first structure includes the stator unit (cf. Supplemental Figure S2A), the joint and the initial 50 nm of the rotor (cf. Supplemental Figure S2B top part). The initial 50 nm of the rotor are extended by 413 nm to its total length of 463 nm by a separate DNA origami nanostructure (cf. Supplemental Figure SS2B bottom part). The stator structure in square lattice geometry uses the first 7236 nt of an 8064 nt long M13mp18 derived scaffold and consists of two sheets which are stacked onto each other rotated by $90°$. The bottom layer of the stator unit is modified with six biotin-modified staples enabling immobilization on PEG-biotin functionalized glass substrates through addition of NeutrAvidin. The remaining 828 nt of the scaffold are used for the joint and the initial 50 nm of the rotor in honeycomb lattice geometry. The rotor extension structure uses a 7249 nt M13mp18 derived scaffold and contains extensions for the attachment of fluorescently labelled strands. Both structures are folded and purified separately.

Additional information about the structure design can be found in [1], where the same structure is employed.

A

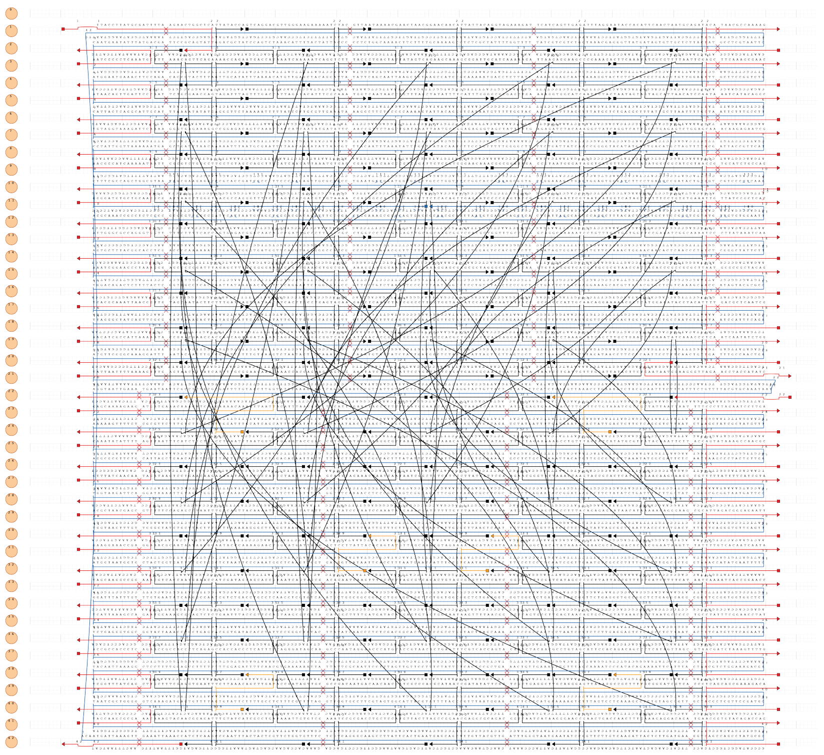

B

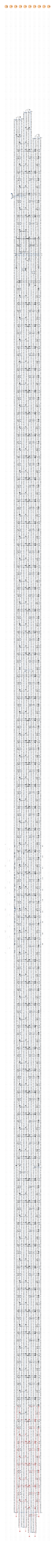

FIG. S2. **Design maps of DNA origami nanostructures** (Figure on preceding page). (a) Cadnano design map of the stator, which uses the first 7236 nt of a 8064 nt scaffold. Biotinylated staples are shown in organge (modification at 5'-end with Biotin and two thymidine bases), passivation staples (red) are extended with four thymine bases at 5'- and 3'-end. (b) Map of the joint and the combined rotor and extension structure. The rotor unit is made from the remaining 828 nt of the 8064 nt scaffold used for the stator. The rotor extension is folded from a 7249 nt long scaffold. Red color indicates the position of staples extended with M1-sequence (taagtgagacccgtacatat) at 5'- and 3'-ends to bind staples modified with fluorescent dyes. Scaffold and staple sequences are included in the design maps. Note: This figure will not print clearly. Some features are only visible in digital format.

#### Estimation of hydrodynamic parameters

As seen in our experiments, the ratcheting behavior of our rotor nanostructures is limited by drag at higher frequencies (>5 Hz). Ratcheting with higher velocities can therefore be accomplished by either decreasing the buffer viscosity or the rotational drag coefficient of the rotor.

*Buffer viscosity.* All experiments were performed in a buffer solution containing 48 % (w/w) sucrose ($\beta$-D-Fructofuranosyl $\alpha$-D-glucopyranoside), which allows to reliably calculate cumulative trajectories because of slowed down rotation of the nanostructures. The increase in buffer viscosity at ambient conditions (T=293.15 K) and therefore the increase in rotational drag can be estimated as $\eta$ = 12.49 $\eta_{H_2O}$ ([5]). A buffer with lower sucrose concentration would allow ratcheting with higher AC field switching frequencies but with the downside of lower observation accuracy.

*Rotational friction coefficient of the rotor.* The rotational friction coefficient $\gamma_r$ of the rotor can be estimated as shown by Vogt et al. [1]:

$$\gamma_r = 8F_r\pi\eta R_e^3 = 1.1 \cdot 10^{-21}\ Nms.$$

The dimensionless friction coefficient $F_r$ corrects for the shape and rotational geometry of the rotor according to Maria Tirado and José Garcia de la Torre [6]:

$$F_r = \frac{2p^2}{9(\ln p - 0.662 + \frac{0.917}{p} - \frac{0.050}{p^2})}.$$

The equivalent radius of a sphere $R_e$ is defined as $R_e = \left(\frac{3}{2p^2}\right)^{1/3} \frac{L}{2}$ with the same volume as the rotor and the aspect ratio $p$ is defined as $p = L/2r$. Reducing the rotor length would therefore be one way to reduce the rotational drag and allow higher rotation velocities.

### SIMULATION OF THE ROTOR MOVEMENT

#### Langevin equation

The movement of the rotor arm within its $2\pi$-periodic potential $E(\phi)$ can be described by the Langevin equation:

$$\gamma_r \dot{\phi} + \frac{dE}{d\phi} = \sqrt{2k_BT\gamma_r} \cdot \eta, \quad (1)$$

where $\gamma_r$ is a rotational friction constant (see previous section for an estimate of $\gamma_r$ for the rotor arm), which is related to the rotational diffusion coefficient via $D_r = k_BT/\gamma_r$. $D_r$ has units of $\mathrm{rad}^2/\mathrm{s}$, and thus $\gamma_r$ has units of $\mathrm{N} \cdot \mathrm{m} \cdot \mathrm{s}/\mathrm{rad}^2$. In the fluctuating term on the RHS, $\eta(t)$ is a Gaussian random variable with

$$\langle \eta(t) \rangle \; = \; 0 \quad (2)$$

$$\langle \eta(t)\eta(t') \rangle \; = \; \delta(t - t') \quad (3)$$

We can rearrange the equation to

$$\dot{\phi} = -\frac{D}{k_BT}\frac{dE}{d\phi} + \sqrt{2D} \cdot \eta, \quad (4)$$

For our numerical simulations, we use the Euler-Maruyama update rule:

$$\phi_n = \phi_{n-1} - \frac{D}{k_BT}\left(\frac{dE}{d\phi}\right)_{n-1} \Delta t + \Delta W, \quad (5)$$

where $\Delta W$ is a Gaussian random variable with zero mean and standard deviation $\sqrt{2D\Delta t}$. For simulations we set the value of the drag term to $\gamma_r = 1.1\ \mathrm{pN} \cdot \mathrm{nm} \cdot \mathrm{s} = 0.275\ k_BT \cdot \mathrm{s}$, which was estimated for a cylindrical rod in sucrose-containing buffer as discussed in the previous section. The simulations were implemented in Python.

#### Potential landscapes

As described in the main text, the rotor's intrinsic potential energy landscape is modeled by

$$E_{in}(\phi) \approx k_BT[A + B\cos 2\phi + C\cos\phi] \quad (6)$$

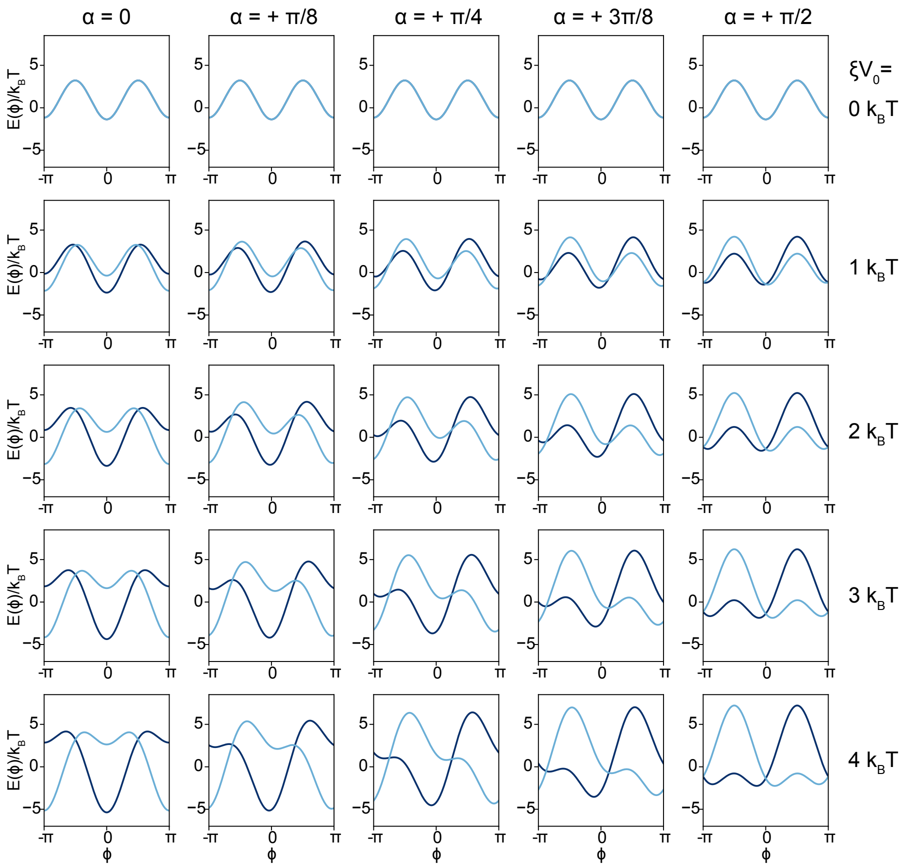


FIG. S3. **Potentials for different angular mismatches at low external fields**. The shapes of the total potential $E(\phi)$ in the two half-cycles of the externally applied square wave voltage are shown (dark blue: negative half-cycle, light blue: positive half-cycle) for different angles $\alpha$ and voltages $\xi V_0$. The potentials are roughly symmetric in the cases $\alpha = 0$ and $\alpha = \pm\pi/2$. The asymmetric change of $E(\phi)$ in the other cases gives rise to a ratchet-like effect that results in a net directed movement in the potential.

with the fit parameters $A = 0.98, B = -2.24, C = -0.108$. The externally applied voltage changes the potential according to

$$E_{ext}(\phi, t) = \xi V_0 \cos(\phi - \alpha) \times \text{sgn}(\sin \omega_0 t), \tag{7}$$

where $\xi$ denotes a factor that converts the externally applied voltage $V_0$ (our experimental control variable) to energy, and $\alpha$ is the angular mismatch between the externally applied field direction

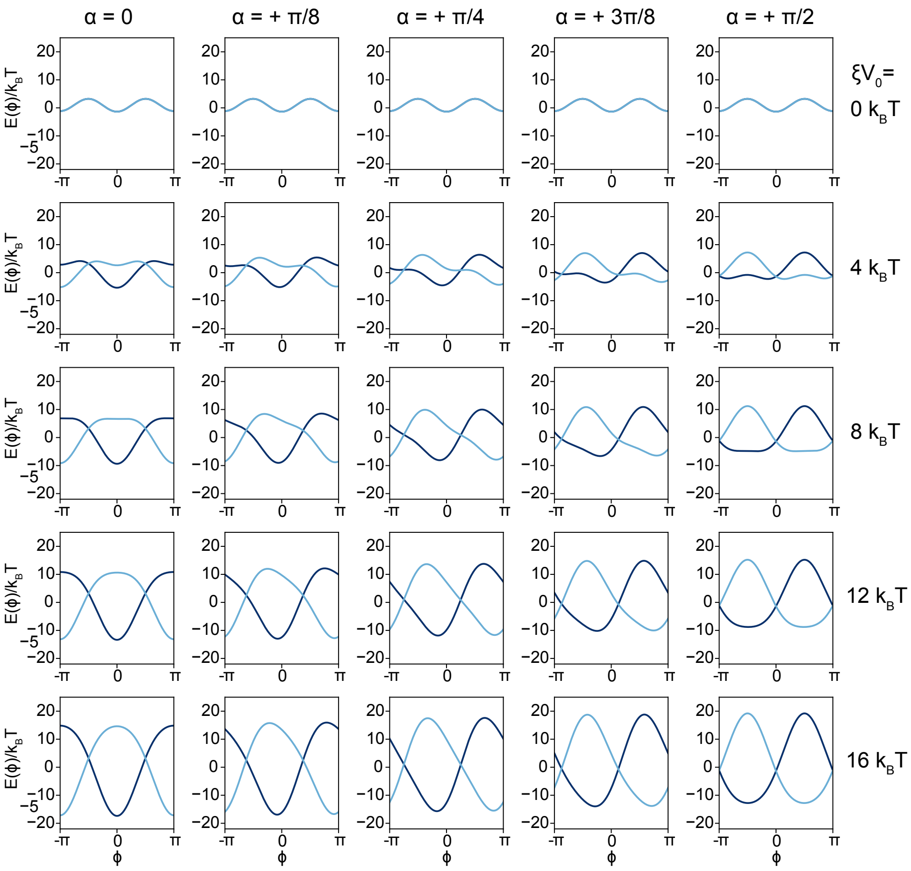


FIG. S4. **Potentials for different angular mismatches at higher external fields**. As in the previous figure the total potential $E(\phi)$ is shown in the two half-cycles of the externally applied square wave voltage (dark blue: negative half-cycle, light blue: positive half-cycle). The potential has two minima in the interval $[-\pi, \pi]$, when $\xi V_0 \lesssim 4.55\ k_B T$, and only one minimum for $\xi V_0 \gtrsim 4.55\ k_B T$.

and the rotor arm platform. The total potential is thus given by:

$$\begin{aligned} E_{tot}(\phi, t) &= E_{in}(\phi) + E_{ext}(\phi, t) \\ &= k_B T[A + B\cos 2\phi + C\cos\phi] + \xi V_0 \cos(\phi - \alpha) \times \mathrm{sgn}(\sin\omega_0 t). \end{aligned} \quad (8)$$

We can also write

$$E_{tot}(\phi, t) = k_B T[A + B\cos 2\phi + C\cos\phi] \pm \xi V_0 \cos(\phi - \alpha), \quad (9)$$

where the '+' sign applies to one half period, and the '−' sign to the other half-period of the periodically switched field. Example plots of $E_{tot}$ for different values of $\alpha$ and $V_0$ are shown in Fig. S3 (low voltage or 'Brownian' regime) and Fig. S4 (high voltage or 'quasi-deterministic' regime).

As stated in the main text, when the external field is increased, the potential minima and maxima that initially were at $\phi \approx 0$ and $\phi \approx \pi/2$, respectively, move towards each other and merge in a saddle point at $\phi = \pi/4$. The saddle point can be numerically calculated by solving $dE_{tot}(\phi)/d\phi = 0$ and $d^2E_{tot}(\phi)/d\phi^2 = 0$ for $\phi$ and $V$, which for $\alpha = -\pi/4$ results in $\xi V_0 \approx 4.55\ k_BT$ (the value slightly differs for different $\alpha$).

**Simulations for low voltages - Brownian regime**

Simulations performed for different values of angular positions at a fixed external voltage in the 'low' regime ($\xi V_0 = 4k_BT$) show the same behavior as seen in the experiments (Fig.S5). The externally applied voltage did not induce any net movement for angular mismatches $\alpha = 0$ or $\alpha = \pm\pi/2$. Movement was CCW ($\omega > 0$)for $\alpha < 0$ and CW ($\omega < 0$) for $\alpha > 0$, with maximum absolute velocities achieved for $\alpha = \pm\pi/4$. A plot of the mean angular velocity against the angle $\alpha$ shows the same sinusoidal dependence $\propto \sin(2\alpha)$ as suggested by the experiments (Fig.S6). In the simulations, we also artificially set the noise term in the Langevin equation to zero ($\Delta W = 0$), which amounts to simply solving the classical overdamped equation of motion in the potential $E_{tot}$. In this case, we observed no motion in the potential except for an initial movement from a random initial angle towards the nearest minimum (see Fig.S5(b)). This behavior shows the necessity for Brownian fluctuations for the generation of movement in this regime, underlining the Brownian motor behavior of the system for low voltages.

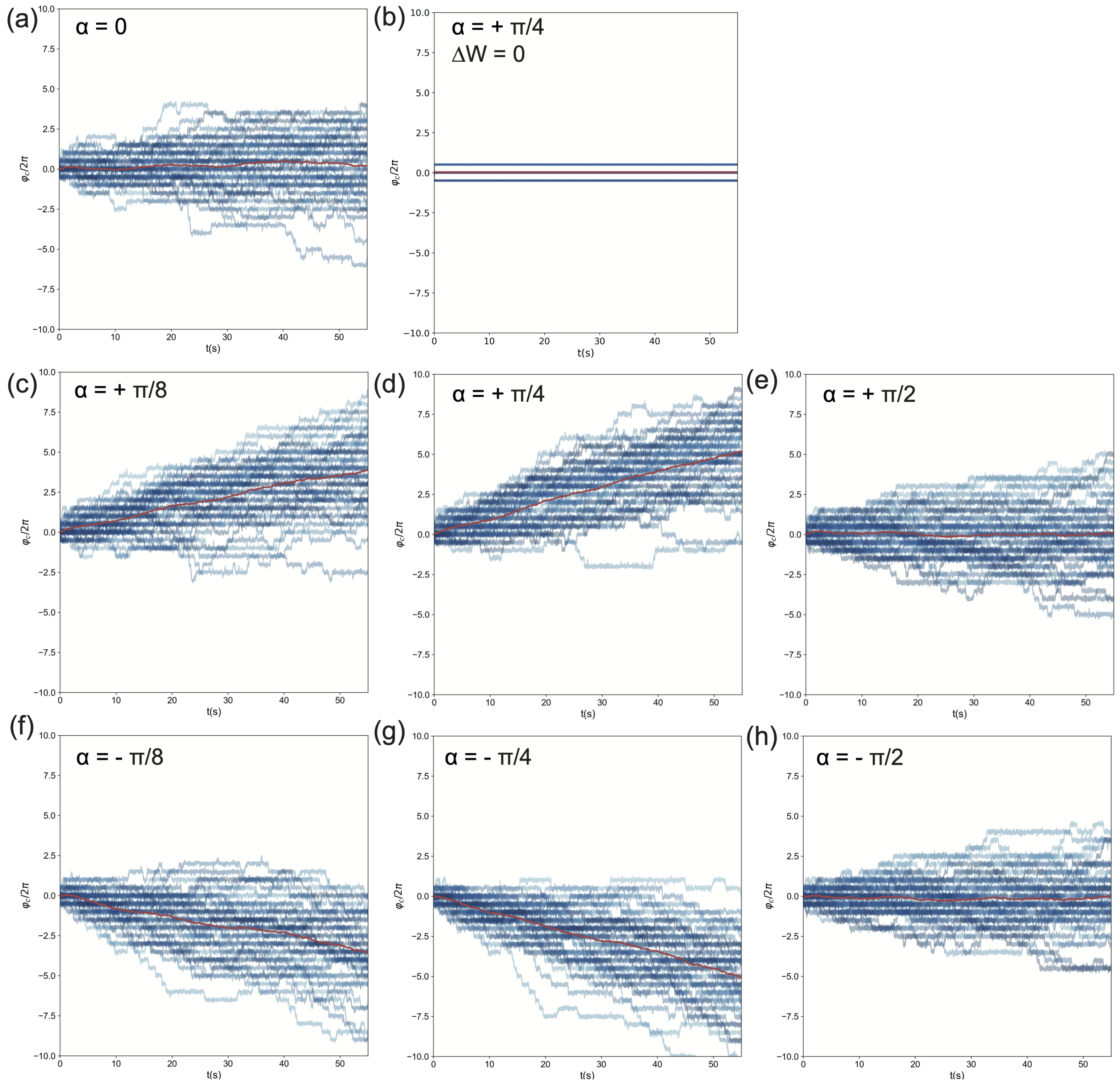


FIG. S5. **Langevin simulations for $\xi V_0 = 4k_BT$.** In the subfigures, 50 simulated trajectories are shown for each of the indicated angular mismatches. The average trajectory is shown as a red line. There is no net directional movement for (a) $\alpha = 0$, and (e,h) $\pm\pi/2$. The movement is CCW for negative $\alpha$ (c,d) and CW for positive $\alpha$ (f,g). In (b), we also show a simulation, where we simply set the fluctuating term $\Delta W = 0$. In this case the system starts at a random position in the interval $[-\pi, \pi]$ and then quickly settles into one of three potential minima (at $\phi = 0, \pm\pi$).

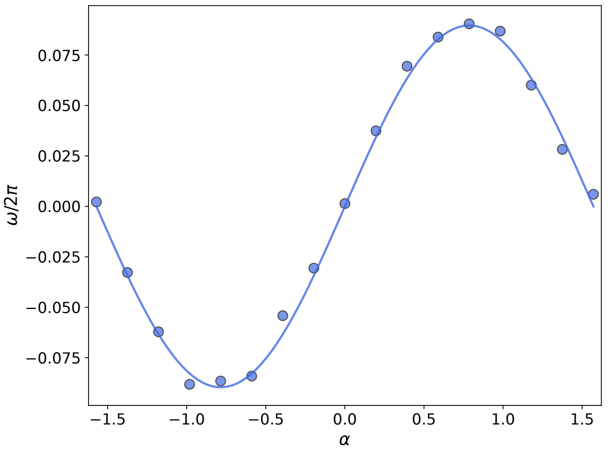


FIG. S6. **Dependence of the angular velocity $\omega$ on the angular mismatch $\alpha$ (simulation).** The data points are the average speeds calculated from 50 Langevin dynamics simulations for $\xi V_0 = 4k_BT$ for each angle. The line plot is a fit of the function $A \sin 2\alpha$ to the data ($A = 0.09$/s), showing that the maximum speeds are attained for $\alpha = \pm\pi/4$.

### Simulations for high voltages

Simulations for higher voltages (Fig. S7) demonstrate the transition from the Brownian to the deterministic regime. As expected, there is no net directional movement for $\alpha = 0$ also in this regime (Fig. S7 (a,d,g,j)). For $\alpha = -\pi/4$ the movement becomes increasingly faster for increasing $\xi V_0$ (Fig. S7 (b,e,h,k)). We also included simulations, in which we set the fluctuation term $\Delta W = 0$ (Fig. S7 (c,f,i,l)). Interestingly, for $\xi V_0 = 8\ k_BT$ this results in no net movement at all, even though at this point the potential already has the shape of a single minimum landscape. For $\xi V_0 = 10\ k_BT$, the $\Delta W = 0$ simulation still results in slower movement than the simulation in the presence of fluctuations, demonstrating that in an intermediate voltage range, thermal fluctuations still support directional movement. Above $\xi V_0 = 12\ k_BT$ this situation changes and, e.g., for $\xi V_0 = 16\ k_BT$ the movement is slower in the presence of fluctuations compared to the simulations for $\Delta W = 0$. In this case the velocity without fluctuations attains its maximum possible speed, where the angular movement of the rotor is precisely clocked by the external frequency, i.e., $\omega/2\pi = f_0 = 5$ Hz. Apparently, for higher voltages the thermal fluctuations lead to occasional idle cycles or even back-steps, which are not present in the purely deterministic case.

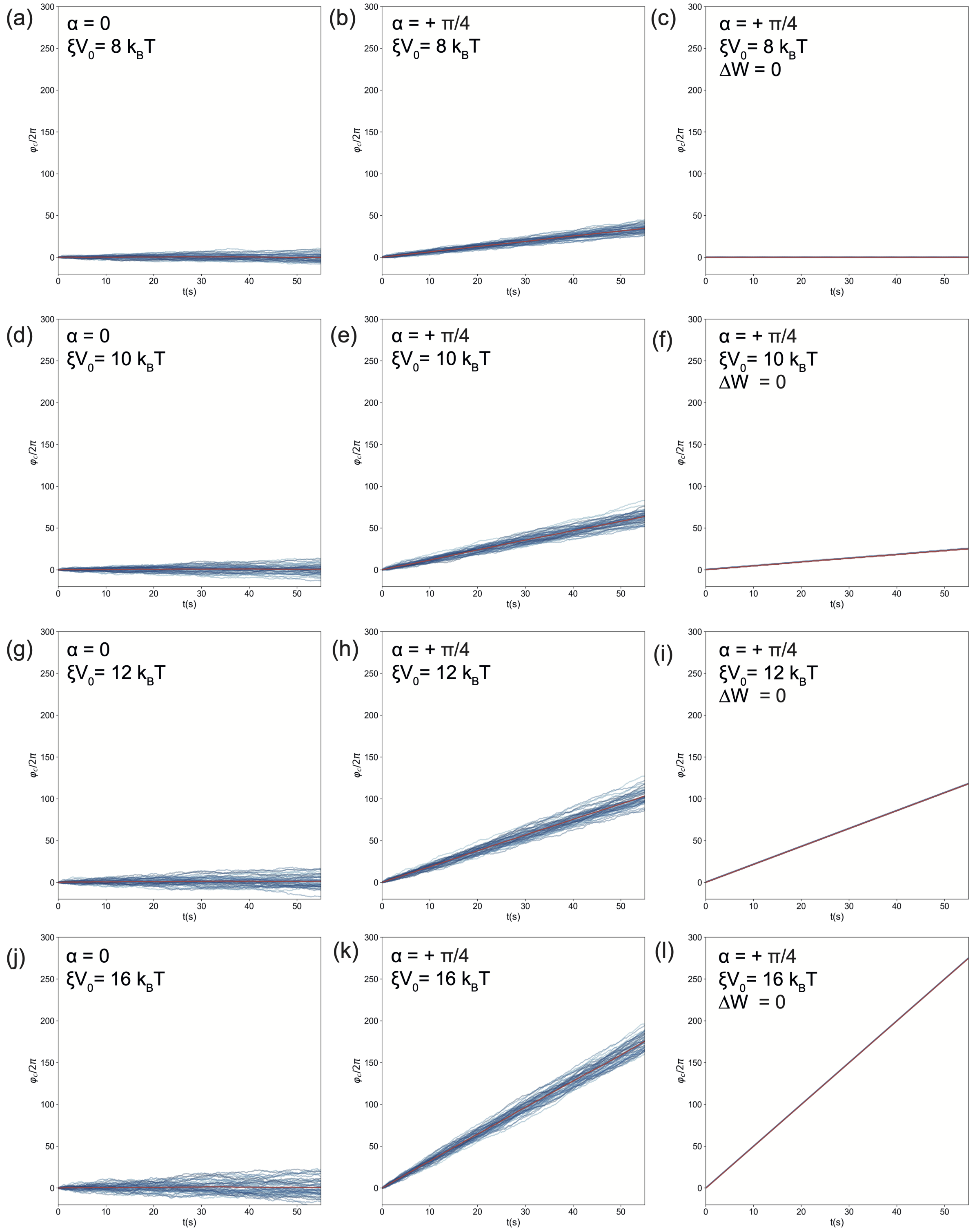


FIG. S7. **Langevin simulations for higher voltages**. (a-c) show simulations for $\xi V_0 = 8 k_B T$, (d-f) corresponds to $V_0 = 10\ k_B T$, (g-i) show $V_0 = 12\ k_B T$, and (j-l) show $V_0 = 16\ k_B T$ (cf. the potential landscapes in Fig.S4.

**Voltage and frequency dependence**

We finally explored the voltage and frequency dependence of the rotor using our simple Langevin model of the system's dynamics. We first performed the simulations for varying $\xi V_0$ for a constant $f_0 = 5$ Hz. As shown in Fig. S8, similar as in the experiment the rotor speed rises steeply with increasing values of the external voltage. The speed saturates at around $\xi V_0 = 20\ k_B T$ and then declines, which is in good qualitative agreement with our experimental observations.

The experimental data shown in Fig. 4(c) of the main text suggests that the maximum speed is achieved for $V_0 \approx 150$ V. Comparison of the experimental data with our simulation suggests a conversion factor of $\xi \approx 0.1\ k_B T/\mathrm{V}$ (see Fig. S8), which is higher than that estimated from previous experiments. However, in the simulations the shape of the voltage dependence is strongly affected by the choice of the hydrodynamic drag $\gamma_r$, where the maximum speed is achieved for smaller $V_0$ with lower values of $\gamma_r$. We also have to note that our simple model most likely does not capture effects that might occur at excessive bias voltages. Further, as noted in the main text, our estimated (intrinsic) potential was obtained by Boltzmann inversion of the angular distribution of the rotor arm, which only has very few data points in the transition regions separating the minima. Therefore, the shape of the underlying potential landscape is not really known, and thus an exact correspondence of simulation and experiment cannot be expected.

Variation of the externally applied frequency for a fixed voltage $V_0$ results in a similar frequency-dependence of the rotor speed as in the experiments (an example for $\xi V_0 = 16 k_B T$ is shown in Fig. S9). Notably, the rotor speed shows a pronounced maximum around $f = 5$ Hz as in the experiment. Simulations show that for high enough voltages ($\xi V_0 \gtrsim 8 k_B T$) the position of the peak varies with voltage roughly linearly (Fig.S11).

The frequency response can be qualitatively understood as the result of the interplay of hydrodynamic drag and external driving. After switching the potential, movement of the rotor arm to the next available minimum has to occur quick enough before the potential is switched back again. Otherwise the arm would return to its initial position, performing an idle cycle. For higher frequencies the time available becomes shorter. On the other hand, stronger tilting of the potential by the external voltage leads to a higher drift velocity, and thus the 'cutoff frequency' becomes higher with voltage. Conversely, the cutoff frequency is reduced for higher drag.

More quantitatively, from the classical equation of motion we obtain as the drift velocity of

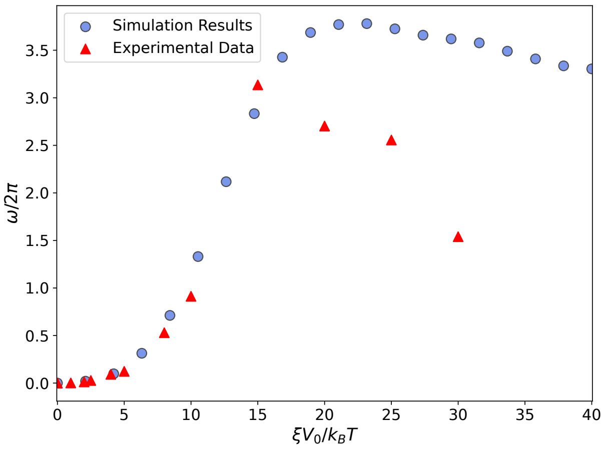


FIG. S8. **Dependence of the rotor arm frequency on the externally applied voltage.** Each point is the average speed obtained from 100 Langevin simulations for the given values of the scaled external voltage $\xi V_0$. For comparison, the experimental data from Fig.4(c) of the main paper are shown as red symbols. For these data, the scaling factor was chosen to be $\xi = 0.1\ k_BT/\mathrm{V}$.

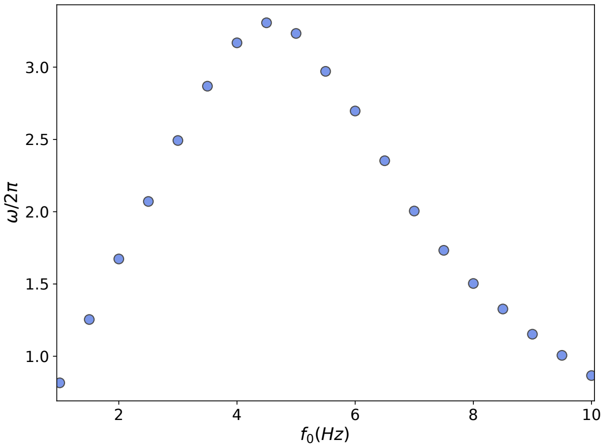


FIG. S9. **Frequency dependence of the rotor arm movement for $\xi V_0 = 16 k_B T$.**

the rotor arm

$$\omega_{drift} = -\frac{1}{\gamma_r}\frac{dE}{d\phi} \tag{10}$$

To make a simple estimate, we can assume that for large external voltages the potential is strongly tilted and the force term can be approximated by the constant

$$-\frac{dE}{d\phi} \approx \frac{2\xi V_0}{\Delta\phi}, \tag{11}$$

where $\Delta\phi$ is the angular distance between the maximum and next minimum of the potential (which turns out to be a little larger than $\pi$ - cf. the plots in Fig. S4). We also assume that the position of the arm after switching is close to the maximum of the switched potential, and now has to reach the next minimum at a distance $\Delta\phi$.

This leads to the requirement:

$$\omega_{drift} \cdot \frac{T}{2} = \omega_{drift} \cdot \frac{1}{2f} \approx \frac{2\xi V_0}{\gamma_r \Delta\phi} \cdot \frac{1}{2f} \gtrsim \Delta\phi, \tag{12}$$

or:

$$f < f_c := \frac{\xi V_0}{\gamma_r \Delta\phi^2} \tag{13}$$

Thus the cutoff frequency $f_c$ is expected to be $\sim V_0/\gamma_r$ for large enough $V_0$.

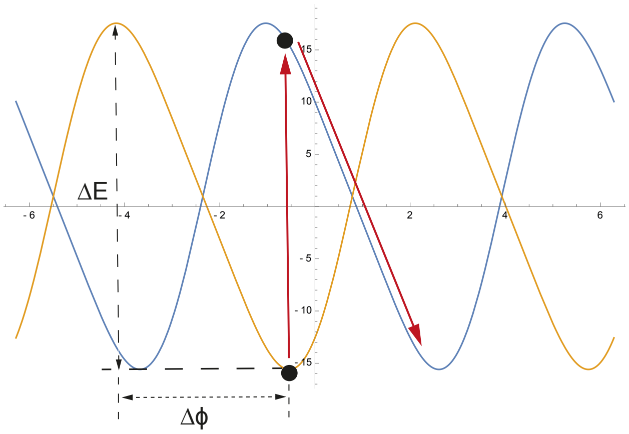


FIG. S10. **Potential switching and frequency dependence.** The rotor arm intially resides in a minumum of the orange landscape. After switching its position is close to a maximum of the blue landscape and then drifts down the potential to the next minimum. For large external voltages, the switching amplitude is given roughly by $\Delta E \approx 2\xi V_0$. The distance between maximum and next minimum is $\Delta\phi$. The average torque is thus given by $2\xi V_0/\Delta\phi$ and the average drift velocity by $\omega_D = 2\xi V_0/(\gamma_r \Delta\phi)$.

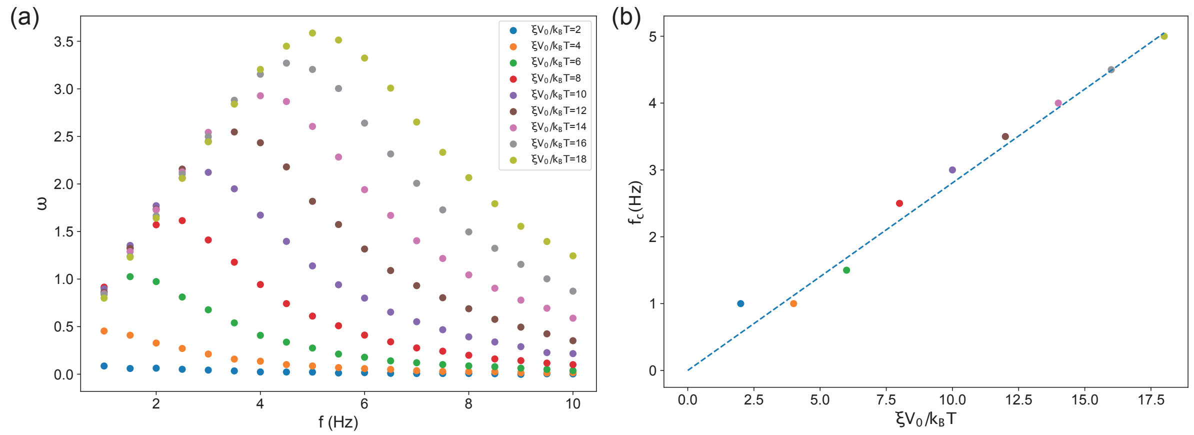


FIG. S11. **Potential switching - frequency and voltage dependence (simulation).** a) The frequency dependence of the angular velocity is shown for different voltages. b) Frequency of the maximum $\omega$ as a function of voltage. The dashed line is given by $f_c = \frac{\xi V_0}{\gamma_r \Delta\phi^2}$ with $\Delta\phi = 3.6$ and $\gamma_r = 0.275\ k_B T \times$ s.

---
